# The GLP-1R Agonist Semaglutide Reduces Motivated Running and Alters Dopamine Dynamics in the Nucleus Accumbens

**DOI:** 10.1101/2025.10.08.681212

**Authors:** Ethan P. Foscue, Joseph R. Trinko, Jaysen Lara Jiménez, Edward Kong, Summer L. Thompson, Kiera Stankewich, Anouk M. Corstens, Mireille J. Serlie, Jane R. Taylor, Ralph J. DiLeone

## Abstract

Glucagon-like peptide-1 receptor (GLP-1R) agonists have recently emerged as powerful tools for the treatment of obesity through their ability to suppress food intake. However, their effects on non-ingestive motivated behaviors remain incompletely understood. Here, we show that the long-acting GLP-1R agonist semaglutide (SG) suppresses voluntary wheel running in both lean and diet-induced obese mice. Importantly, this suppression of activity was not caused by hypophagia and was accompanied by decreased motivation, with SG-treated mice displaying reduced effort for wheel access in a progressive ratio task. Real-time measurements of dopamine via fiber photometry revealed specific dopamine changes in the nucleus accumbens at both the beginning and end of running bouts, with SG-treated animals showing amplified dopamine dynamics at these key behavioral timepoints. Collectively, these data reveal important non-ingestive behavioral effects of GLP-1R agonism and suggest a role for dopamine circuits in mediating reductions of volitional activity following SG treatment.

## Introduction

Glucagon-like peptide-1 receptor (GLP-1R) agonists, originally developed for type 2 diabetes mellitus, have now emerged as among the most effective pharmacological tools for the treatment of obesity. The therapeutic effect is primarily driven by reductions in food intake, with stabilized, or long-acting, agonists having shown the greatest potency for inducing weight loss (Rubino et al., 2021; Wilding et al., 2021). These GLP-1–based agents significantly improve health outcomes, in part by reducing risks for cardiometabolic and obesity-related comorbidities (Lincoff et al., 2023). Long-acting GLP-1R agonists have been taken by over 12% of adults in the U.S., with over 30 million individuals projected to receive prescriptions within the next 5 years (Montero et al., 2024 KFF Health Tracking Poll). The widespread use of these therapeutics demands that we have a more complete understanding of their mechanisms and broader behavioral effects, which might provide clearer guidance for future use.

While long-acting GLP-1R agonists likely function via multiple brain regions, reward circuits are likely to mediate any changes in behavior. Animal models have provided critical insights into the mechanisms by which GLP-1R agonists reduce ingestive behavior. In rodents, GLP-1 agonists have been shown to reduce the consumption of highly palatable foods and attenuate operant responding for food rewards, partly through effects on mesolimbic dopamine circuits (Alhadeff et al., 2012; Alhadeff and Grill, 2014; Dickson et al., 2012; Dossat et al., 2013, 2011; Fortin and Roitman, 2017; Hsu et al., 2018, Hsu et al., 2015; Konanur et al., 2020; López-Ferreras et al., 2019; Maske et al., 2018; Mietlicki-Baase et al., 2014, 2013; Wang et al., 2015). Consistent with these findings, clinical studies have revealed that GLP-1R agonists modulate activity in reward- and motivation-related brain regions, reducing their responsivity to food cues while promoting reductions in food cravings and overall caloric intake (Bae et al., 2020; Blundell et al., 2017; Farr et al., 2016; Schlögl et al., 2013; ten Kulve et al., 2015; van Bloemendaal et al., 2015, 2014; Wilding et al., 2021).

Emerging evidence suggests that GLP-1R agonists also influence broader reward-related behaviors beyond reduction in food intake. GLP-1R agonists reduce operant responding for non-food rewards, such as alcohol or drugs of abuse (Egecioglu et al., 2013c, Egecioglu et al., 2013a, Egecioglu et al., 2013b; Erreger et al., 2012; Herman et al., 2023; Reddy et al., 2016; Schmidt et al., 2016; Shirazi et al., 2013; Sørensen et al., 2015; Vallöf et al., 2016), highlighting the potential use of GLP-1R agonists in addressing a range of maladaptive behaviors associated with dysregulated reward systems. Emerging data from human studies and clinical observations also support a potential application for curbing addictive behaviors, such as alcohol or nicotine use (Farokhnia et al., 2025; Hendershot et al., 2025; Jerlhag, 2025; O’Keefe et al., 2025; Qeadan et al., 2025). These findings raise the critical question of how long-acting GLP-1R agonists influence other non-ingestive, reward-related behaviors.

Exercise is a compelling domain in which to probe the broader effects of GLP-1R agonists, as it engages reward and motivation systems while serving as a major contributor to energy balance and overall physical and mental health (Ding et al., 2025; Kyu et al., 2016; Liu et al., 2022; Noetel et al., 2024; Singh et al., 2025; Wahid et al., 2016). Physical activity relies on and engages neural pathways that influence motivation, overlapping with the systems targeted by GLP-1R agonists (Ekkekakis and Zenko, 2024; Ling et al., 2024; Roefs et al., 2023; Stillman and Erickson, 2019; Stults-Kolehmainen et al., 2023). Voluntary wheel running, a well-validated proxy for exercise in rodents, provides a robust framework to evaluate GLP-1R agonist effects on physical activity, as it is intrinsically rewarding and engages neural systems regulating motivation and energy expenditure (Greenwood et al., 2011; Greenwood and Fleshner, 2019; Rhodes et al., 2003; Skovbjerg et al., 2024). Here we present data evaluating the effects of the GLP-1R agonist semaglutide (SG) on running behavior, the motivation to run, and modulation of associated dopamine dynamics in mice.

## Methods

### Animals

Animal experiments were carried out in accordance with Yale University School of Medicine Institutional Animal Care and Use Committee (IACUC) regulations. All protocols were approved by the Yale IACUC, and all surgeries were performed in compliance with approved pre- and post-operative care. C57BL/6J young adult male and female mice were purchased from the Jackson Laboratory (Bar Harbor, ME), at approximately 8-10 weeks of age (for animal numbers for each experiment, see Supplemental Table 5). Prior to testing, mice were group housed on a normal light-dark cycle (7am-7pm) and placed on either a standard rodent chow diet (Teklad Global, Inotiv 2918), or a high-fat diet (Western Diet, TestDiet 5TJN) for 5 weeks prior to being single housed with the running wheel.

Fiber photometry animals were single housed on a reverse light-dark cycle (7am-7pm) following surgery and kept on standard rodent chow for the duration of the experiment.

### Drugs

Semaglutide (“SG”; Biosynth, FS171058) was diluted in 100% DMSO at 1 mg/ml and then aliquoted and frozen. Preparation for injection consisted of thawing the aliquots at 37°C and brief sonication to ensure SG was fully solubilized. The final injection solution was 24 µg/ml of SG and 2.4% DMSO in saline, and animals were injected with 60 µg/kg/day (s.c.). Pair-fed (PF) and vehicle animals received 2.4% DMSO in saline. Drug and vehicle were delivered daily at 3pm, four hours before the onset of the dark cycle, for all experiments.

### Apparatus

For free running experiments, animals were singly housed in cages with vertical metal wheels (Allentown LLC, Allentown, NJ). Wheel rotations were mechanically counted using micro switches with a hinge lever and were recorded using ClockLab 3 and ClockLab 3 Analysis (Actimetrics, Lafayette, IN).

Operant experiments took place in the above home cages with running wheels. Two nosepokes with cue lights (Med Associates, Fairfax, VT) were then added to these cages, along with a brake built from a modified operant lever for rats (Med Associates, Fairfax, VT). This brake would extend out, blocking the path of the wheel and locking it in place. Mice could then nosepoke on various schedules to unlock the wheel and gain the ability to run. Behavioral testing was controlled using Med-PC 5 software (Med Associates, Fairfax, VT).

Fiber photometry experiments took place in an open topped rodent cage (6” x 11” x 8”) containing a saucer style running wheel (EveryYay Exercise Saucer for Small Animals, Small, 3891347) to prevent the wheel from interfering with the tether. The wheels were present in the home cages for the duration of the experiment, to familiarize mice with the apparatus.

### Behavior

Mice were initially trained in the wheel cages for two weeks to allow them to establish familiarity with the wheel and a baseline level of running. During the initial free running experiment mice remained on their standard diet, and were divided into two groups, receiving either SG or vehicle injections. Pair feeding with vehicle injection was attempted, but significant food wastage by SG animals made this difficult to achieve in the first group of animals. The experiment was subsequently rerun using only a SG and pair-fed group. During the free running experiment with mice receiving high fat diet, the animals were split into the same SG and vehicle injected groups, with vehicle mice also having their food intake matched to SG mice. Food intake was matched by weighing the food in the hopper once each day, and pair feeding was accomplished by providing the pair-fed group with the mean weight of food the SG group had consumed on the previous day.

Accurate pair feeding was achieved by providing SG-injected animals with approximately 10 grams of food per day, with the bedding of the cages monitored for signs of food wastage. For seven days, mice with SG or vehicle at 3pm and wheel counts were binned in 24-hour periods beginning at 3pm.

For progressive ratio experiments, following the initial free wheel training described above, mice were first trained to nosepoke on a fixed ratio (FR) 1 schedule for food pellets (F0071, BioServ, Flemington, NJ). They were then left in their wheel cages overnight on a FR1 schedule for wheel access, where a single nosepoke response would unlock their wheel for ten minutes. One nosepoke was deemed the active port as indicated by an illuminated cue light, whereas the other nosepoke served as an inactive control with no programmed consequences. All animals successfully learned the task, without any food restriction, and after five days began progressive ratio testing. Mice were run on an escalating semi-linear reinforcement schedule (Figure 4A) in which each reward (10 minutes of wheel access) required progressively more nosepoke responses than the previous reward. The program began at the start of the dark cycle and ran until it was manually turned off the following day at 3pm, and the breakpoint was defined as the last reward achieved before the animal did not nosepoke for 30 consecutive minutes. The experimental timeline began with five consecutive days of overnight progressive ratio testing with no injection, followed by five consecutive days of overnight PR testing after being injected with SG as described above.

To investigate changes in running on smaller timescales, the recording software was updated to bin wheel turns each second. Using custom Python scripts, binned wheel turns were then compiled into a timeseries for each animal across each day. Exercise behavior was separated into “bouts,” defined by a minimum number of turns and a maximum number of seconds in between turns. For all analyses, these criteria were a minimum of 2 turns and a maximum of 2 seconds in between turns (Figure 5A). Bouts were analyzed in terms of length in seconds, speed in turns per second, and total number of bouts per day.

### Fiber Photometry

The fiber photometry setup consisted of a custom-built set up as previously described (Bond et al., 2020; de Jong et al., 2024; Trinko et al., 2024). Briefly, an LED driver drove LED cubes of 430nm and 490nm, which were passed through 440nm and 490nm bandpass filters, respectively (all parts from Thorlabs, Newton NJ). Light was selectively passed through a 495nm dichroic mirror and filtered through a 520nm bandpass filter (all from IDEX Health and Science, LLC, West Henrietta NY). This signal was captured using a Grasshopper3 2.3 camera (Teledyne FLIR, Charlotte NC) at a rate of 10Hz. Behavioral data (wheel turns) were measured and digitally signaled using a custom Arduino Uno system measuring turns using magnets attached to the wheel and a latching hall sensor. The LED driver, camera, and behavioral signals were driven using a CB-68LP terminal block (National Instruments, Austin TX), and all instruments were controlled using custom built MATLAB code.

pAAV-hsyn-GRAB-DA2m with a titer of 2.4×10^13^ was purchased from Addgene (Watertown, MA). Mice were anesthetized with isoflurane and injected with 0.5 µl of GRAB-DA2m virus unilaterally in the nucleus accumbens (stereotaxic coordinates A/P 1.2, M/L 0.6, D/V −4.5 mm from bregma). A fiber optic cannula (400 µm dia. fiber core, N.A. 0.48) was lowered to the same A/P and M/L coordinates, and D/V −4.2 mm from bregma), and then cemented in place. Two weeks were allowed for recovery and viral expression following surgery and prior to behavioral experiments.

Mice were habituated to the running setup during daily sessions without photometry recording to familiarize them with running while tethered. On recording days, fibers were photobleached for approximately one hour before recordings to minimize background autofluorescence. During behavioral recordings, mice were allowed to run freely for approximately two hours, during which activity was recorded from the signal (490nm) and reference (430nm) streams. Wheel turns were recorded and timestamped into the fiber photometry signal file. For SG testing, mice were injected daily at 3pm for three days prior to the beginning of fiber photometry recordings with SG, and injections continued daily throughout testing, following the conclusion of the subject’s fiber photometry recording session.

### Statistics and Data Analysis

Wheel turn data were compiled using ClockLab 3 Analysis (Actimetrics, Lafayette, IN). MEDPC data was compiled using custom code written in Python, and statistical analysis was performed using the R lme4, coxme, and emmeans packages. For free running experiments, a linear mixed-effects regression was fit to the data, and a type-III ANOVA was performed on the coeVicients to test for main effects and interactions. For multiple comparisons, marginal means for conditions were estimated from the regressions, and compared using t-tests. Specific tests and parameters are reported in supplemental materials. Figures were compiled in GraphPad Prism. For comparisons including baseline effects, baseline was defined as the mean of the last 3 days prior to the beginning of injection. As all experiments included mice of both sexes, sex was included as a covariate in all analyses, and animals were pooled across sex where there was no significant effect.

Fiber photometry data was processed using custom Python scripts and statistically analyzed using the fastFMM R library (Loewinger et al. 2025). Briefly, raw photometry signal was filtered using a Butterworth bandpass filter with 2Hz and 0.001Hz cutoffs, before it was regressed onto reference signal to eliminate potential artifacts. It was then z-scored across the entire session. For event related analysis, bouts were identified using similar criteria as above but instead using a minimum of 10 turns and a 5 second interbout interval, to ensure bout-related events were well spaced.

Baseline signal was collected by randomly sampling signal windows of the noted length from throughout the session. These signal windows were then labeled with necessary trial information and compiled into tables for functional linear mixed modelling (FLMM) analysis.

## Results

### Semaglutide reduces free running distance in lean mice

We first tested the effects of SG in lean animals in a voluntary running model. Mice were given free access to running wheels for 14 days, during which the running distance increased to over 10km per day (Supplemental Figure 1). The mice were then treated with 60 µg/kg SG daily to test the effects of acute and chronic treatment on running distance, and a linear mixed-effects regression (LMER) was fit to their performance. As expected, mice receiving SG experienced an immediate and significant weight loss, which continued throughout the injection period (Figure 1A). Mice receiving SG ran significantly less distance than their baseline (−17093 ± 970 wheel turns, t=17.624, p<0.0001) and vehicle injected controls (−15090 ± 3350 wheel turns, t=4.500, p=0.0017) (Figure 1B; Supplemental Table 1 for multiple comparisons). To further examine the effects of repeated injections of SG over multiple days, the total distance run on each day was compared to the mean baseline running. The number of wheel turns for SG animals was significantly reduced during each treatment day, relative to the vehicle injected controls (Figure 1C; Supplemental Table 1).

**Figure 1.**
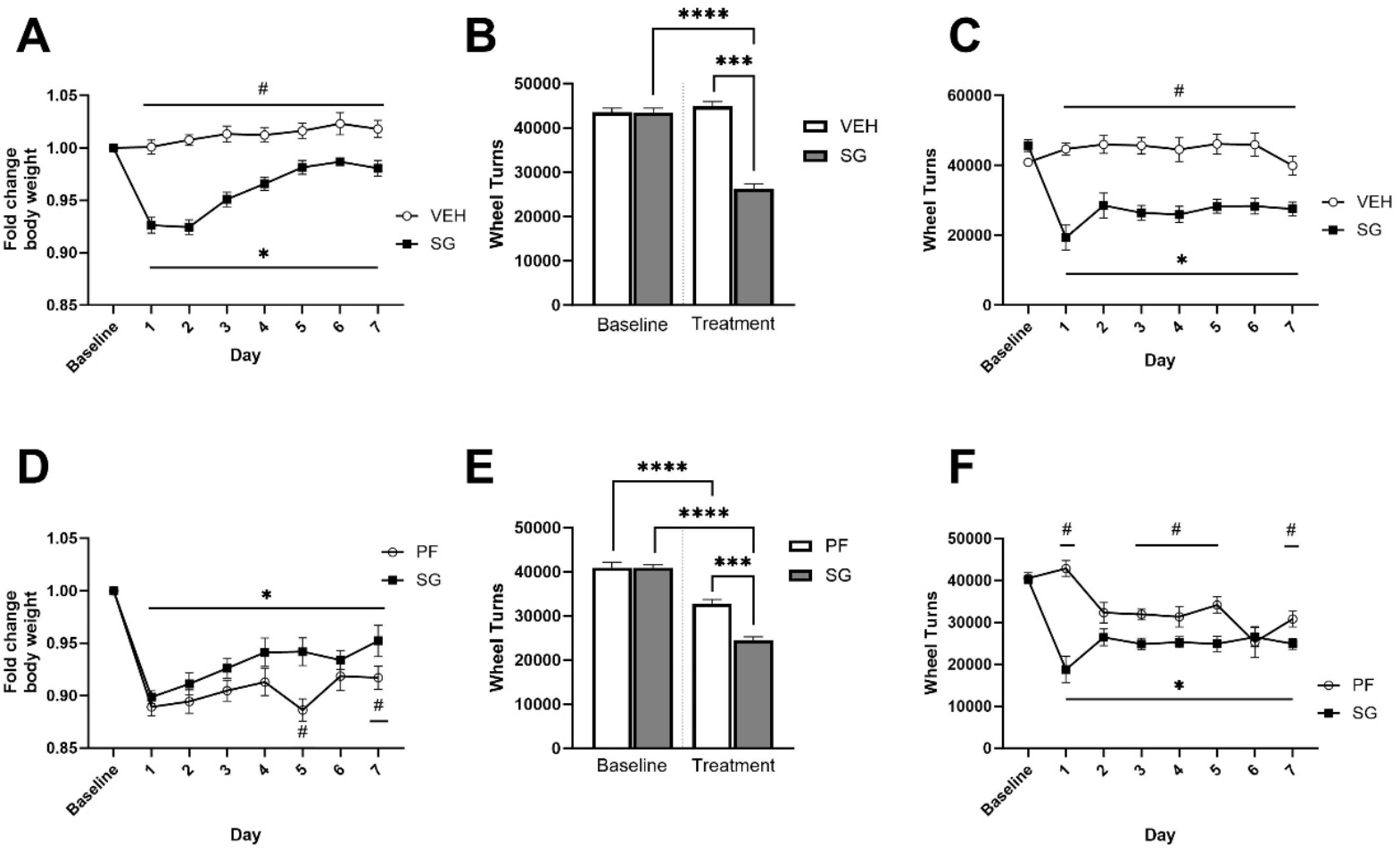
Semaglutide (SG) reduced free running distance in lean animals. See supplemental table 1 for specific statistical results (A) Fold change from each animal’s baseline (defined as the 3 days before treatment) weight during SG injection in lean animals. SG significantly reduced animals’ body weight at all days of treatment relative to baseline (*) and vehicle injected controls (#) (B) Mean wheel turns during baseline and SG injection in lean animals. SG significantly reduced wheel turns in lean mice relative to the animal’s baseline levels (p<0.0001, n=11) and vehicle injected controls (p<0.0001, n=11). (C) Mean daily wheel turns at baseline and during each individual day of injection in lean animals. SG significantly reduced wheel turns relative to the animals’ baseline (*) and relative to vehicle injected controls (#) at all days of treatment, with no differences at baseline. (D) Fold change from each animal’s baseline weight during SG injection and pair-fed (PF) in lean animals. (E) Mean wheel turns during baseline and treatment of SG injected animals and pair-fed controls. SG (p<0.0001, n=8) and pair feeding (p<0.0001, n=8) both significantly reduced wheel turns during treatment, but SG injected animals ran significantly less than pair-fed animals (p=0.0005, n=16). (F) Mean daily wheel turns at baseline and during each individual day of SG injections or pair-fed in lean animals. SG significantly reduced wheel turns relative to the animals’ baseline (*) and relative to pair-fed controls (#) at days 1, 3, 4, 5, and 7 of treatment, with no differences at baseline. All error bars represent S.E.M.

To test whether reduced activity was caused by reduced caloric intake, we repeated the experiment with pair-fed vehicle mice. Pair-fed mice had their food intake matched to the SG group (Supp.Figure 2) and showed similar weight loss (Figure 1D). Pair-fed animals also ran less than at baseline (−7844 ± 1320 wheel turns, t=5.922, p<0.0001), yet more than SG animals (8382 ± 1870 turns, 4.489, p=0.0005) (Figure 1E). Analysis of daily running revealed that SG mice ran less than pair-fed mice during most of the treatment, and less than their baseline on all days (Figure 1F; Supplemental Table 1). These data demonstrate the acute and sustained effects of SG on running distance beyond the effects of caloric reduction alone.

### Semaglutide reduces free running distance in diet-induced obese (DIO) mice

To extend the analysis to a more clinically relevant condition, we next tested the effects of SG after 5 weeks of high-fat diet. DIO mice receiving SG ran significantly less during treatment than baseline (−12527 ± 1600 wheel turns, t=7.842, p<0.0001) and vehicle injected pair-fed controls (−11171 ± 3300, t=3.389, p=0.0055) (Figure 2A), with significant per-day reductions relative to baseline (Figure 2B). SG-treated mice also ran significantly less than pair-fed controls on days 1 and 2 of SG injection (Figure 2B). To address the possibility that the difference in running across conditions was primarily driven by the reduction on day 1 of SG treatment, a separate model was fitted with SG day 1 removed, and difference between groups remained (−8583 ± 3680 wheel turns, t=2.335, p=0.0353). As expected, SG-injected mice also reduced their food intake (−29.7% ± 4.9%, t=6.022, p<0.0001) and body weight during the injection period, and pair-fed mice displayed similar body weight loss (Figure 2C, and 2D). Direct comparison of the free running distance of chow and DIO groups revealed no significant difference, either at baseline or during any of the days of SG injection (Figures 3A and 3B). As expected, DIO mice lost a higher percentage of their body weight than lean animals (5.16% ± 0.833%, t=6.193, p<0.0001; Figure 3C).

**Figure 2.**
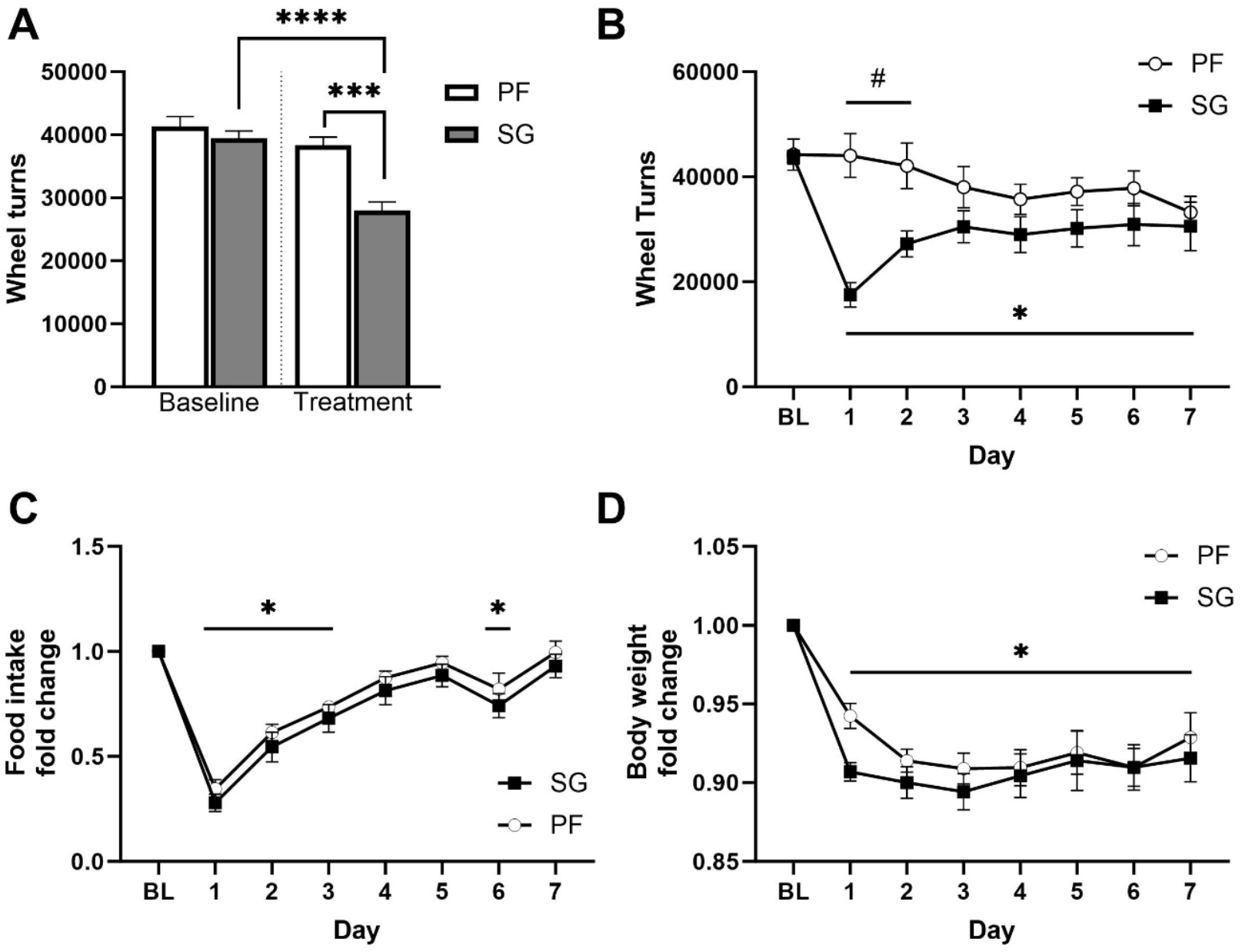
Semaglutide (SG) reduced free running distance in diet-induced obese (DIO) mice. See supplemental table 2 for specific statistical results. (A) Mean wheel turns during baseline (defined as the 3 days before treatment) and SG injection in DIO animals. SG significantly reduced wheel turns relative to the animals’ baseline levels (p<0.0001) and vehicle injected, pair-fed (PF) controls (p<0.001). (B) Mean daily wheel turns at baseline and during each individual day of injection in DIO animals. SG significantly reduced wheel turns relative to the animals’ baseline (*) on all days of treatment and relative to vehicle injected controls (#) on the first two days of treatment, with no differences at baseline. (C) Percentage of each animal’s baseline food intake during SG injection in DIO animals. SG significantly reduced food intake relative to baseline at days 1, 2, 3 and 6 of injection. Pair-fed animals received matched food during the injection period. (D) Percentage of each animal’s baseline weight during SG injection in DIO animals. SG significantly reduced animals’ body weight at all days of treatment relative to baseline (*), while there were no significant differences between SG injected and pair-fed, vehicle injected mice on any day of treatment. All error bars represent S.E.M.

**Figure 3.**
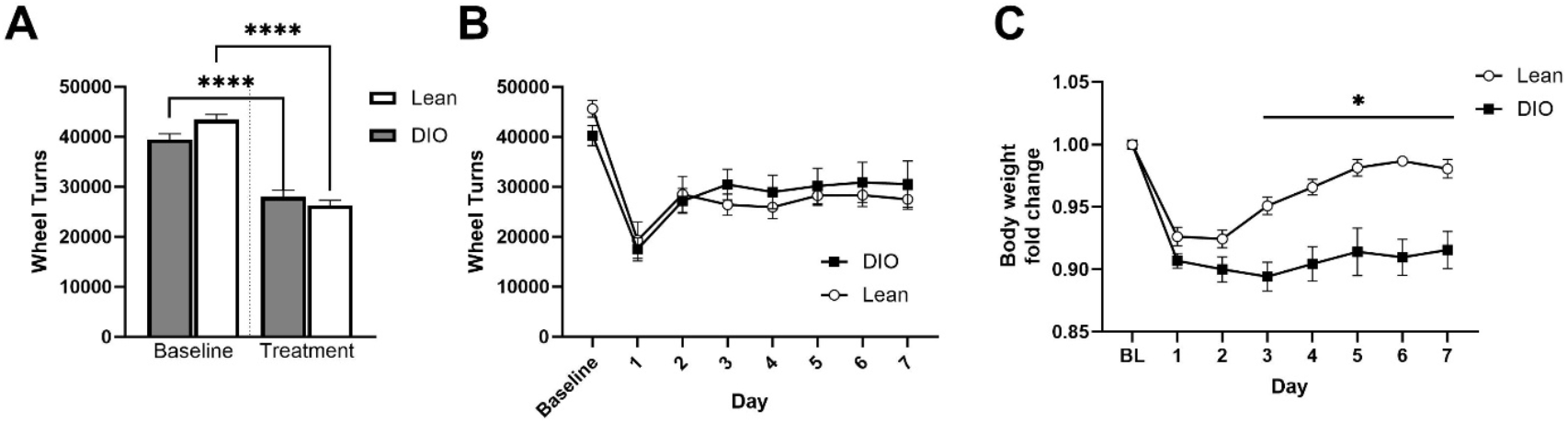
Diet had no differential effect on free running distance in semaglutide (SG) injected animals. See supplemental table 3 for specific statistical results. (A) Mean wheel turns during baseline (defined as the final 3 days of treatment) and SG injection in lean and DIO animals. There were no significant differences in wheel turns between the two diets at baseline (p=0.3782) or during SG injection (p=0.5683). (B) Mean daily wheel turns at baseline and during each individual day of injection in lean and DIO animals injected with SG. There were no significant differences in wheel turns between the two diets at any day of injection. (C) Percentage of each animal’s baseline weight during SG injection in lean and DIO animals. Lean animals body weight relative to baseline was significantly higher than that of DIO animals on days 3-7 of injection (*). All error bars represent S.E.M.

### Semaglutide administration reduces progressive ratio performance in an operant wheel running task

To assess motivation to access wheels, mice in both diet groups were injected with SG and tested on a progressive ratio (PR) task (Figure 4A). In both diet groups, SG administration significantly reduced the number of wheel accesses achieved in a Cox proportional hazards analysis (Lean: BL/SG ratio=0.214 ± 0.0423, z=-7.788, p<0.0001, DIO: BL/SG ratio=0.478 ± 0.0821, z=-4.299, p<0.0001). There was no significant difference between the diets at baseline (p=0.5220) and during SG administration (p=0.2208, Figure 4B). Notably, the largest reduction in performance occurred on the first day of SG treatment, likely reflecting nonspecific malaise, such as transient gastrointestinal effects (Figure 4C). To address this issue, data were reanalyzed excluding the first day of both baseline and SG phases, and the SG-induced reduction in PR performance remained robust in both diet groups (Lean: BL/SG ratio=0.166 ± 0.0394, z=-7.561, p<0.0001, DIO: BL/SG ratio=0.416 ± 0.0818, z=-4.463, p<0.0001).

**Figure 4.**
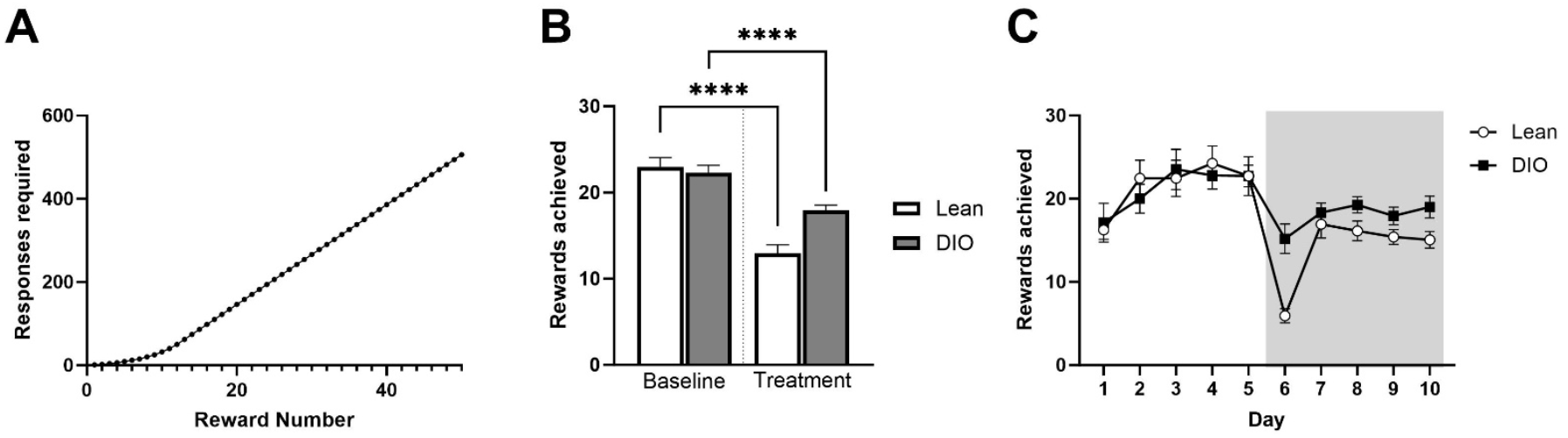
Semaglutide (SG) administration reduces progressive ratio (PR) performance in an operant wheel running task. (A) Progressive ratio schedule during the task, displaying number of nosepokes required for a 10 minute wheel access reward (B) Mean rewards achieved during baseline and SG injection in a PR task in lean and DIO mice. There were no significant differences in rewards achieved between the two diets at baseline (p=0.4952) or during SG injection (p=0.2533). Animals achieved significantly fewer rewards during SG injection than they did during baseline (p<0.0001 for both diets). (C) Mean daily rewards achieved during baseline and SG injection. All error bars represent S.E.M.

### Semaglutide changes running bout structure

To investigate how SG changes the behavioral structure of mouse running on shorter timescales, 32 lean mice (16 males and 16 females) were trained for 14 days, and their running bouts were collected and analyzed based on running bout criteria (Figure 5A-C). During training, the bout structure of animals changed significantly (Figure 5D-F). Notably, the bout count per day significantly decreased across training sessions (Liner mixed-effects regression, LMER, β=-45.07, SE=4.48, t(30.22)=-10.06, p<0.0001, CI[-54.39, −36.15], n=526 sessions, Figure 5D). Bout length in seconds increased significantly across training sessions (LMER, β =2.51, SE=0.19, t(28.74)=13.21, p<0.0001, CI[2.13, 2.90], n=526 sessions, Figure 5E). Bout speed also significantly increased across training sessions (LMER, β=0.027, SE=0.002, t(27.13)=17.47, p<0.0001, CI[0.0240, 0.030], n=526 sessions, Figure 5F). Given these training-related adaptations, only the final three days of training were used as the within-subject baseline for evaluating SG effects, as detailed below.

**Figure 5.**
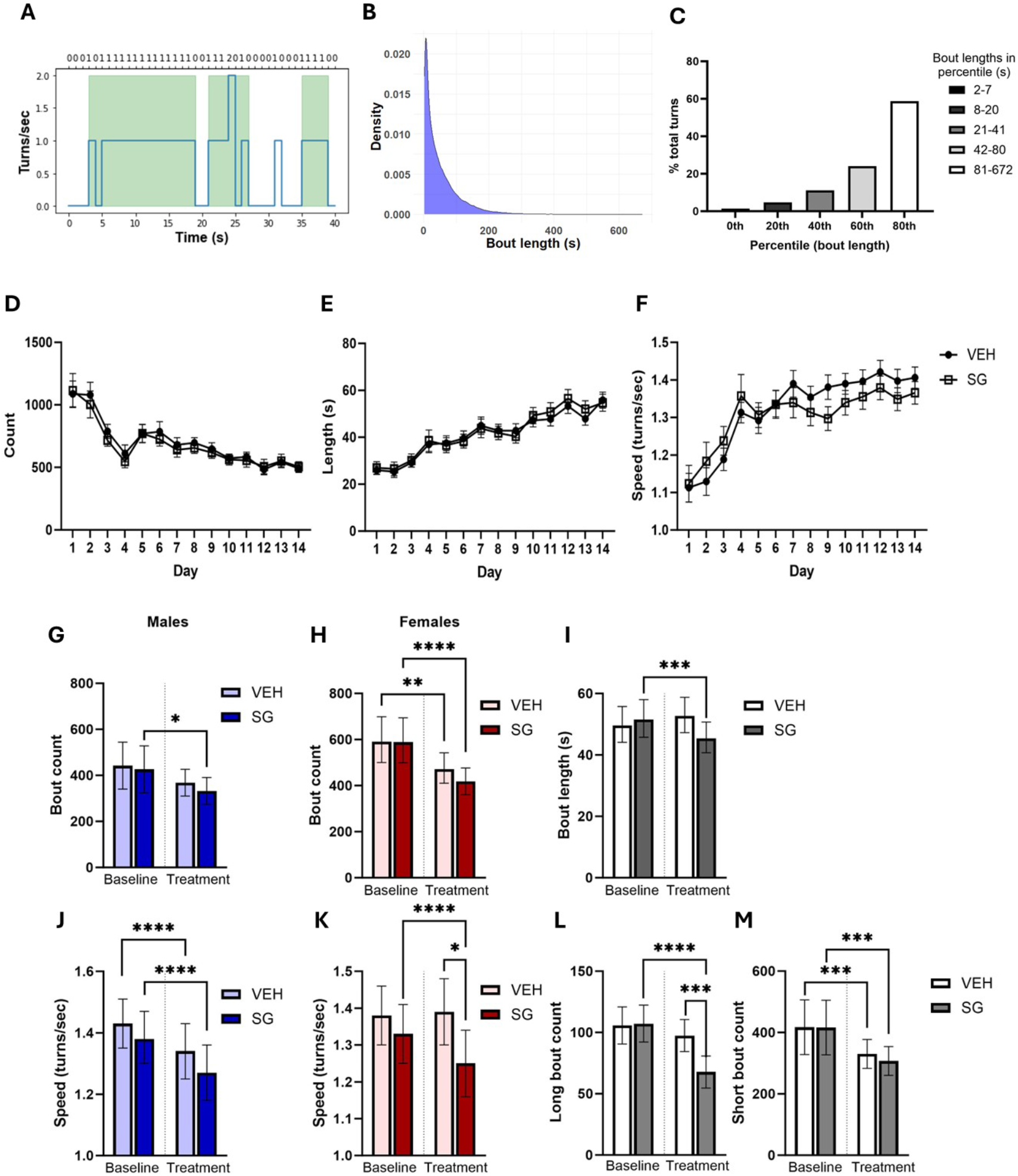
Semaglutide (SG) administration modifies running bout structure. Bouts were compiled and analyzed using the criteria of 2 minimum turns and 2 maximum interturn interval. (A) Representative examples of the data readout and bout classification of running behavior. Bottom axis represents seconds, and top displays turns counted by the software. Green shaded areas represent time periods grouped as a bout, with this example depicting three bouts based on our criteria and thus excluding the single wheel turn. (B) Kernel density estimation plot of bout length during the last three days of training, showing the right skewed distribution of bout length, indicating most bouts are relatively short. (C) Bouts were binned into percentiles based on length, and the percent of total turns that occurred in each bin is displayed, indicating the most wheel turns occur within longer bouts. (D-F) Bout count, mean bout length, and mean bout speed during each day of training. Bout count significantly decreased across all days of training (p<0.0001, n=526 sessions), while mean bout length (p<0.0001, n=526 sessions) and mean bout speed (p<0.0001, n=526 sessions) both significantly increased across training, All error bars represent S.E.M. (G-H) Bout count per day was higher in females across all conditions (p= 0.011, n=315 sessions), and significantly decreased from training to test in both SG and vehicle injected animals (p=0.002, n=315 sessions), but there was no significant interaction between phase and drug. Baseline refers to the last 3 days of training, and Test refers to all days of injection (I) Bout length in seconds significantly decreased in SG treated but not VEH treated animals during testing (p=0.0004, n= 138,039 bouts, pooled sexes) All error bars represent 95% CI. (J-K) Bout speed in turns/sec during the test phase was significantly decreased relative to baseline in VEH injected males (p<0.0001, n=20,593 bouts), SG injected males (p<0.0001, n=27,844 bouts), and SG injected females (p<0.0001, n=38,310 bouts) but not in vehicle injected females. All error bars represent 95% CI. (L-M) SG administration’s effect on the number of bouts over 81s (80^th^ percentile of training bouts) and under 81s. There was a significant reduction in longer bout counts in SG injected animals (p<0.0001, n=315 sessions, pooled sexes) but not vehicle injected control mice. Both SG and vehicle injected had significantly fewer test phase bouts under 81s than during baseline (p=0.005, n=315 sessions) but there was no difference between the groups during injection. All error bars represent 95% C.I.

### Semaglutide administration changes the behavioral structure of wheel running

Repeated SG administration differentially affected specific phases of the bout structure. An LMER on log-transformed total bout number per day revealed a main effect of sex (p = 0.011, n = 315 sessions), with males exhibiting fewer bouts than females across all conditions (Figure 5G–H). A significant main effect of phase (training vs. test) was also observed (p = 0.00191,n = 315), but no significant interaction with drug treatment (p = 0.09), suggesting that SG did not differentially affect overall bout number (see Supplemental Table 4 for detailed statistics for all data in this section).

To assess how SG impacted bout length, data were fitted to a generalized linear mixed model. No main effect of sex was detected (p = 0.751), and including sex did not improve model fit; therefore, data were pooled across sexes (Figure 5I). A significant interaction between phase and drug emerged (p = 0.0004, n = 138,039 bouts). SG-injected animals showed significantly shorter bouts during the test phase compared to their training baseline (p = 0.0024, n = 66,152 bouts), whereas vehicle-injected control mice showed a non-significant trend toward increased bout length (p = 0.063, n = 72,124 bouts).

In order to assess how bout speed was affected by SG, an LMER was fitted to the animal’s turns per second during bouts. There was a significant interaction between phase and drug (p = 0.0022, n = 138039), and between phase and sex (p = 0.001, n = 138039). These effects were driven by significant decreases in bout speed during SG treatment in both sexes: SG-injected males (p < 0.0001, n = 27,844 bouts), SG-injected females (p < 0.0001, n = 38,309 bouts), and vehicle-injected males (p < 0.0001, n = 31,204 bouts). No significant change was observed in vehicle-injected female mice (Figure 5J–K).

To further explore the selective impact of SG on bout structure, we separated bouts by duration using the 80th percentile of the baseline distribution (81 seconds) as a cutoff for longer bouts. An LMER was then used to assess changes in the number of short (<81 s) and long (>81 s) bouts per day (Figure 5L-M, pooled sexes). For short bouts, a significant main effect of phase was observed (p = 0.005, n =315), indicating a general decline across both groups. In contrast, for long bouts, a significant interaction between phase and drug treatment emerged (p < 0.0001, n = 315). SG-injected mice completed significantly fewer long bouts post-treatment compared to baseline (p < 0.0001, n = 157 sessions), and significantly fewer than vehicle-injected controls during the test phase (p = 0.003, n = 222 sessions).

### Opposing dynamics in nucleus accumbens extracellular dopamine around the start and end of free running bouts

Mice were allowed to engage in voluntary free running while extracellular dopamine was measured in the nucleus accumbens using the GRAB-DA sensor(Sun et al., 2020). Signal dynamics at the beginning and end of running bouts were analyzed using functional linear mixed modeling (FLMM) to assess how fluorescence traces differed between bout-aligned time windows and randomly selected control epochs sampled throughout the session. Bouts were defined using a criterion of ≥10 wheel turns and ≥5 s interbout interval. FLMM revealed a significant reduction in the estimated mean dopamine signal relative to random signal windows from approximately −6 to +5 seconds relative to bout onset (95% CI, Figure 6A), with the decrease occurring ∼1 second prior to the first wheel turn. At bout termination, the estimated signal mean showed a transient increase from approximately 0 to +2.5 seconds following the final wheel turn. This increase was followed by a rapid decline, with reductions in dopamine signal observed from approximately +5 to +10 seconds post-bout completion (95% CI, Figure 6B).

**Figure 6:**
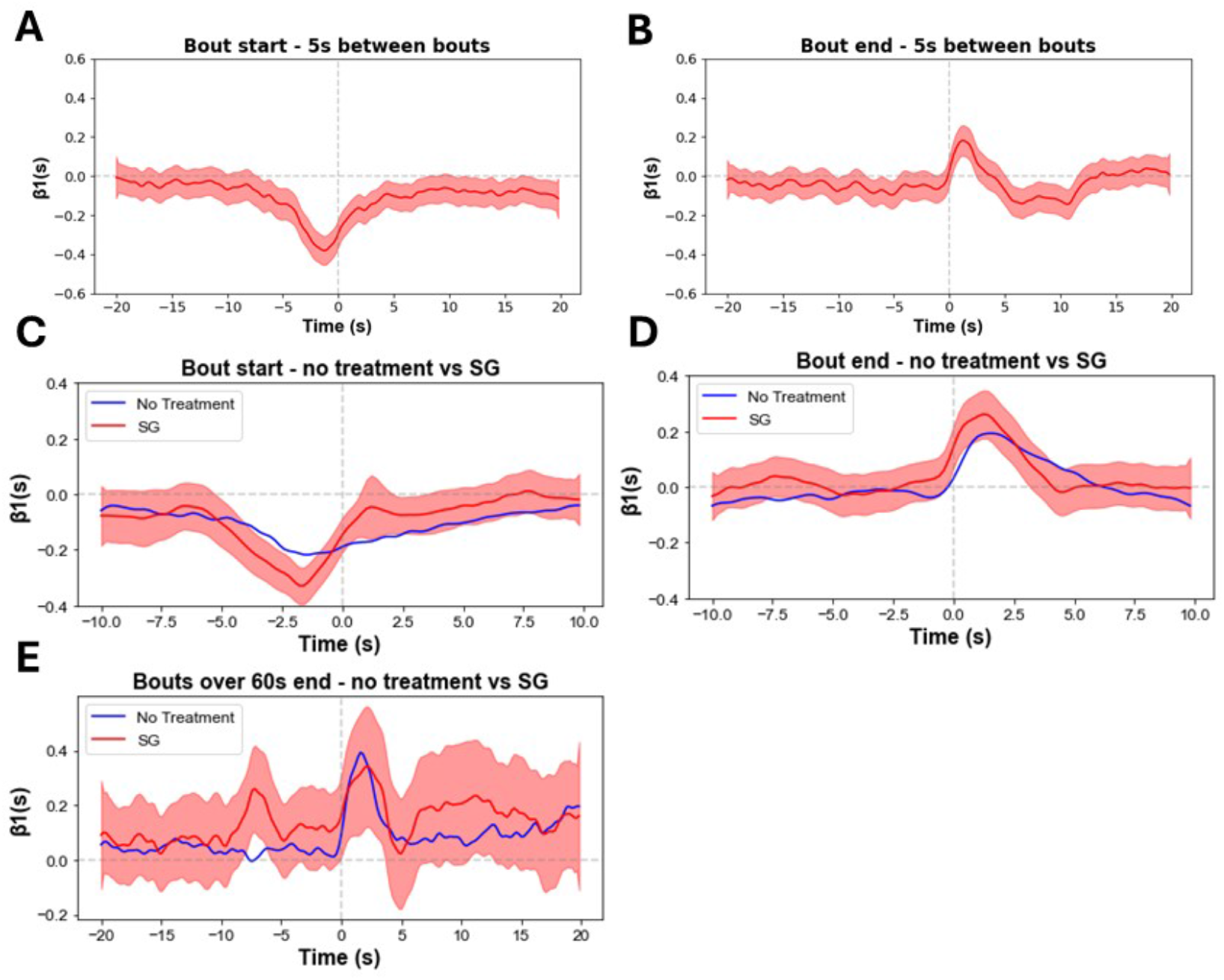
Semaglutide (SG) administration alters dopamine dynamics in the nucleus accumbens during free running. GRABDA was used to measure extracellular dopamine in the NAc in free running animals, with signal locked to bout events (bouts were defined as a minimum of 10 turns and 5s interbout interval except where noted). (A-E) FLMM Estimate of functional coeVicients at each timepoint around bout events centered at 0s (line) with joint 95% confidence intervals (shaded). Areas in which the origin (x=0) is not included in the shading indicate a significant difference in mean signal relative to random signal windows. (A) Animals displayed a significant decrease in signal from ∼[-5, 5] seconds around the start of a running bout (n=1812 bouts). (B) The same analysis in the interval [-20, 20] seconds around the end of free running bouts revealed a significant increase in signal from ∼[0, 2.5] seconds, which then decreased below random signal levels from ∼[5, 10] seconds (n=1811 bouts). (C-D) Estimated effect of SG injection on mean signal magnitude relative to no treatment in the interval [-10, 10] around the start and end of bouts. Areas in which the red shaded area (SG) does not overlap the blue line (no treatment reference) indicate a significant change in mean signal magnitude during treatment (n=3300 bouts). (E) As SG treatment significantly reduced the number of longer bouts during overnight sessions, the end of bouts longer than 60s were isolated during fiber photometry sessions and compared between no treatment and SG. FLMM analysis revealed an increase in estimated mean signal from ∼[-8, −6] seconds prior to bout end during SG relative to no treatment (n=679 bouts).

### Semaglutide administration amplifies dopamine dynamics at start and end of free running bouts

To investigate how repeated SG administration affected dopamine dynamics in free running animals, mice received 3 days of SG injection without recording. Starting on day 4, mice continued to receive SG daily and were again monitored using fiber photometry during free running as above. FLMM analysis revealed effects of SG on extracellular dopamine dynamics at bout onset compared to no-treatment sessions from approximately −4 to −1 seconds relative to the first wheel turn (n = 3,300 bouts; Figure 6C). A subsequent transient increase in dopamine signal was observed from approximately +1 to +2.5 seconds after bout onset relative to untreated bouts (n = 3,300). Bout termination was analyzed similarly and revealed a small but significant elevation in dopamine signal from ∼0 to +1 second following the final wheel turn during SG treatment (Figure 6D). This was followed by a brief reduction in estimated mean signal between +4.6 and +5.0 seconds post-bout (Figure 6D).

These data indicate that dopamine dynamics at the start and end of running bouts are preserved in temporal structure but exhibited amplified magnitude shifts following SG treatment. Given that SG administration reduced average bout duration in overnight recordings, we next isolated bouts exceeding 60 seconds during fiber photometry sessions to determine whether bout length modulated dopamine dynamics. In this subset, FLMM analysis revealed a unique significant enhancement in mean signal during SG treatment from approximately −7.6 to −5.6 seconds prior to bout termination, relative to no-treatment controls (n = 679 bouts; Figure 6E).

## Discussion

The present study demonstrates that the long-acting GLP-1R agonist semaglutide suppresses voluntary wheel running behavior in both lean and DIO mice, revealing a previously underappreciated influence of GLP-1 signaling on non-ingestive motivated behaviors. While GLP-1R agonism is well characterized in the context of central energy balance regulation and limbic reward pathway modulation (Alhadeff et al., 2012; Alhadeff and Grill, 2014; Dickson et al., 2012; Dossat et al., 2011; Fortin and Roitman, 2017; Konanur et al., 2020; López-Ferreras et al., 2019, 2018; Mietlicki-Baase et al., 2013), its effects on volitional behaviors such as exercise are largely unexplored. Our results suggest that semaglutide reduces physical activity not through energetic impairment, but by attenuating behavioral motivation, likely via overlapping mechanisms that also mediate food seeking and reward valuation. These findings underscore the importance of considering GLP-1R agonist effects on motivated activity and provide a mechanistic framework for how GLP-1R signaling interacts with shared reward circuits underlying both ingestive and non-ingestive actions.

A central observation of this study is that SG reduced both the quantity of spontaneous running and the willingness of animals to work for wheel access in a progressive ratio task, suggesting a reduction in exercise-driven motivation. Importantly, these effects on free running were observed in both lean and obese mice and persisted under conditions where food intake was controlled (i.e., pair feeding), pointing to direct effects on motivational processes, rather than reductions that are secondary to hypophagia or weight loss. These results are consistent with prior studies showing that GLP-1R activation diminishes operant responding for diverse classes of reinforcers, including palatable foods, alcohol, and psychostimulants (Aranäs et al., 2023; Egecioglu et al., 2013c; Schmidt et al., 2016; Sørensen et al., 2015; Vallöf et al., 2016). Given the shared neurocircuitry governing food reward, substance use, and voluntary exercise (DiLeone et al., 2012; Ruiz-Tejada et al., 2022; Stults-Kolehmainen et al., 2020), it is plausible that SG diminishes the reinforcing value of running by modulating common dopaminergic circuits, such as the nucleus accumbens.

While we did not directly manipulate GLP-1 receptors or dopamine signaling in this study, prior work has shown that activation of central GLP-1R can attenuate dopamine release in the nucleus accumbens (Fortin and Roitman, 2017; Konanur et al., 2020; Merkel et al., 2025) and modulate synaptic input into dopaminergic neurons (Mietlicki-Baase et al., 2013; Wang et al., 2015). In parallel, voluntary wheel running is known to induce reward-like plasticity in mesolimbic circuits and engage dopamine-associated pathways (Greenwood et al., 2011). Our photometric findings further support this link, revealing temporally structured dopamine fluctuations surrounding running bouts (e.g. pre-bout suppression and post-bout rebound) that are amplified by semaglutide. This pattern is broadly consistent with a model in which semaglutide alters reward prediction errors or the valuation of behavioral outcomes, thereby reducing the anticipatory salience or hedonic impact of voluntary movement.

An alternative, non-mutually exclusive interpretation is that semaglutide suppresses exercise by interfering with the habitual or compulsive components of running behavior. Voluntary wheel running in rodents is supported by both motivational and automatic processes, and can display features reminiscent of compulsivity or automaticity (Eikelboom and Mills, 1988; Novak et al., 2012). In clinical populations, especially anorexia nervosa, excessive/compulsive exercise (hyperactivity) is well documented and closely linked to eating-disorder psychopathology (Davis and Claridge, 1998; Klein et al., 2004; Lichtenstein et al., 2017). Given that habitual and compulsive behaviors are mediated by dorsal striatal circuitry and dopaminergic signaling (Everitt and Robbins, 2013; Graybiel, 2015), the possibility arises that GLP-1R activation dampens habitual behavioral drive by altering striatal circuit dynamics and balance. Future studies using additional behavioral analysis and circuit-specific interventions will be needed to dissociate motivational and habitual contributions to GLP-1–induced reductions in physical activity.

The effects of GLP-1R agonists add to the growing literature defining mechanisms of central regulation of movement and exercise. Voluntary wheel running is intrinsically rewarding for rodents and robustly engages mesolimbic dopamine, hypothalamic, and cortical circuits regulating motivation and energy expenditure (Bastioli et al., 2022; Greenwood et al., 2011; Greenwood and Fleshner, 2019; He et al., 2018; Rhodes et al., 2003; Skovbjerg et al., 2024). Recent studies have supported functional roles for hypothalamic and mesolimbic circuits in regulating voluntary activity in mice (Hwang et al., 2023; Nishitani et al., 2025; Tesmer et al., 2024), and provided evidence of systemic states converging on dopamine circuits to influence exercise (Dohnalová et al., 2022). Together, these findings highlight that voluntary activity is shaped by an interplay of hypothalamic and mesolimbic pathways, underscoring how peripheral signals and pharmacological interventions can modulate dopaminergic substrates of exercise and motivated behavior.

Several limitations of the present work should be considered. First, while wheel running and progressive ratio paradigms are widely used as proxies for exercise motivation, they may not fully capture the multidimensional nature of physical activity in humans across diverse contexts.

Second, although pair feeding controls help dissociate reduced running from caloric restriction, we cannot fully exclude the possibility that energetic or peripheral factors — such as altered muscle function — contribute to the observed suppression of activity. Emerging clinical reports suggest that GLP-1 agonists may induce reductions in lean mass, including skeletal muscle, potentially compromising exercise capacity independently of central effects (Karakasis et al., 2025; Linge et al., 2024; Lu et al., 2025; Neeland et al., 2024; Prado et al., 2024). The extent to which these changes reflect GLP-1R pharmacologically mediated events versus changes that are secondary to weight loss remains unclear.

The present findings have important translational implications. Given the widespread and increasing use of SG in the treatment of obesity and type 2 diabetes (Watanabe et al., 2024), it is critical to understand the full behavioral repertoire influenced by GLP-1R agonism. If SG and related therapeutics blunt voluntary physical activity via motivational suppression, dampened reward processing, or attenuation of habitual exercise, this could have unintended consequences for long-term energy expenditure, weight maintenance, and preservation of fat-free mass. Future clinical studies incorporating objective measures of physical activity (e.g., accelerometry, VO_2_max, or performance tasks) will be essential to determine whether these effects translate to humans, particularly in populations attempting to integrate structured exercise into weight loss or maintenance regimens.

## Acknowledgements

This work was funded in part by the State of Connecticut, Department of Mental Health and Addiction Services, but this publication does not express the views of the Department of Mental Health and Addiction Services or the state of Connecticut. The work was also in part supported by the Charles B. G. Murphy funds (J.R.T.). S.L.T was supported by K99AA029454 and E.K. was supported by the Rosenfeld Science Scholar and the College Dean’s Fellowships from Yale University. We would also like to thank Cathy Duman, Johannes de Jong, Michael Siniscalchi, and Aakash Basu for help and suggestions with the running wheel systems, fiber photometry, and data analysis.

## Notes

### Competing Interest Statement

The authors have declared no competing interest.

