## Supplemental Table 4 for "The GLP-1R Agonist Semaglutide Reduces Motivated Running and Alters Dopamine Dynamics in the Nucleus Accumbens"

**Supplemental Figure – Bout analysis detailed statistics**

Figure 5D – Bout count per day during training

Linear mixed effect regression (LMER), β=-45.07, SE=4.48, t(30.22)=-10.06, p<0.0001, CI[-54.39 -36.15], n=526 sessions

Figure 5E – Bout length per day during training

LMER, β =2.51, SE=0.19, t(28.74)=13.21, p<0.0001, CI[2.13, 2.90], n=526 sessions

Figure 5F – bout speed per day during training

LMER, β=0.027, SE=0.002, t(27.13)=17.47, p<0.0001, CI[0.0240, 0.030], n=526 sessions

Figure 5G-H – Bout count per day during treatment

Main effect of sex: LMER, log transform, β = 0.731, t(27.78) = −2.72, p = 0.011, 95% CI [0.589, 0.910], n = 315 sessions

Main effect of phase: LMER, log transform, β = 0.797, t(29.24) = −3.411, p = 0.00191, 95% CI [0.702, 0.904], n=315 sessions

Interaction between drug and phase: LMER, log transform, β = 0.850, t(28.36) = -1.754, p = 0.09016, 95% CI [0.713, 1.014], n=315 sessions

Figure 5I – Bout length during treatment

Interaction between drug and phase: GLMER, Gamma family, log link, β = 0.849, t = −3.54, p = 0.0004, 95% CI [−0.254, −0.073], n = 138,039 bouts

Estimated marginal mean comparison – GLMER, Gamma family, log link, response scale SG baseline vs treatment: 52.8 s, 95% CI [47.6, 58.6] vs. 46.4 s, 95% CI [41.7, 51.7]; z = 3.038, p = 0.0024, n = 66,152 bouts

Estimated marginal mean comparison – GLMER, Gamma family, log link, response scale VEH baseline vs treatment: 49.7 s, 95% CI [44.4, 55.5] vs. 51.4 s, 95% CI [45.8, 57.8]; z = −1.862, p = 0.063, n = 72,124 bouts

Figure 5J-K – Bout speed during treatment

Interaction between drug and phase: LMER, β = −0.090, SE = 0.027, t(27.85) = −3.371, p = 0.0022, 95% CI [−0.141, −0.040], n = 138,039 bouts

Interaction between phase and sex: LMER, β = −0.100, SE = 0.027, t(27.98) = −3.748, p = 0.001, 95% CI [−0.151, −0.050], n = 138,039 bouts

Estimated marginal mean comparison – LMER

SG injected males baseline vs treatment: 1.38 turns/sec, 95% CI [1.30, 1.47] vs. 1.27 turns/sec, 95% CI [1.18, 1.36]; z = 5.922, p < 0.0001, n = 27,844 bouts

SG injected females baseline vs treatment: 1.33, 95% CI [1.25, 1.41] vs. 1.25, 95% CI [1.16, 1.34]; z = 4.194, p < 0.0001, n = 38,309 bouts

VEH injected males baseline vs treatment: 1.43, 95% CI [1.35, 1.51] vs. 1.34, 95% CI [1.24, 1.43]; z = 4.722, p < 0.0001, n = 31,204 bouts

Figure 5L – Long bout count per day during treatment

Interaction between drug and phase: LMER, β = −31.227, SE = 6.357, t(30.37) = −4.912, p < 0.0001, 95% CI [−43.662, −18.791], n = 315 sessions

Estimated marginal mean comparison – LMER

SG injected mice, baseline vs treatment: 107.2 bouts, 95% CI [92.2, 122.3] vs. 67.8 bouts, 95% CI [54.7, 80.8]; t(29.6) = 8.821, p < 0.0001, n = 157 sessions

Treatment phase, VEH vs SG mice: 97.4 bouts, 95% CI [84.4, 110.5] vs. 67.8 bouts, 95% CI [54.7, 80.8]; t(30) = 3.279, p = 0.003, n = 222 sessions

Figure 5M – Short bout count per day during treatment

Main effect of phase: LMER, β = −87.407, SE = 28.955, t(30.55) = −3.019, p = 0.005, 95% CI [−143.987, −30.780], n = 315 sessions
